# Selecting a new electron transfer pathway for nitrogen fixation uncovers an electron bifurcating-like enzyme involved in anaerobic aromatic compound degradation

**DOI:** 10.1101/2022.10.07.511188

**Authors:** Nathan M. Lewis, Abigail Sarne, Kathryn R. Fixen

**Affiliations:** BioTechnology Institute and Department of Plant and Microbial Biology, University of Minnesota, St. Paul, MN

**Keywords:** nitrogenase, *Rhodopseudomonas palustris*, ferredoxin, NAD^+^-dependent ferredoxin:NADPH oxidoreductase

## Abstract

Nitrogenase is the key enzyme involved in nitrogen fixation and uses low potential electrons delivered by ferredoxin (Fd) or flavodoxin (Fld) to reduce dinitrogen gas (N_2_) to produce ammonia and hydrogen. Although the phototrophic alphaproteobacterium *Rhodopseudomonas palustris* encodes multiple proteins that can reduce Fd, the FixABCX complex is the only one shown to support nitrogen fixation, and *R. palustris* Fix^-^ mutants grow poorly in nitrogen-fixing conditions. To investigate how native electron transfer chains (ETCs) can be redirected towards nitrogen fixation, we leveraged the strong selective pressure of nitrogen limitation to isolate a suppressor of *R. palustris* Δ*fixC* that grows under nitrogen-fixing conditions. We found two mutations were required to restore growth under nitrogen-fixing conditions in the absence of functional FixABCX. One mutation was in the gene encoding the primary Fd involved in nitrogen fixation, *fer1*, and the other mutation was in *aadN*, which encodes a homolog of NAD^+^-dependent Fd:NADPH oxidoreductase (Nfn). We present evidence that AadN plays a role in electron transfer to benzoyl-CoA reductase, the key enzyme involved in anaerobic aromatic compound degradation. Our data support a model where the ETC for anaerobic aromatic compound degradation was re-purposed to support nitrogen fixation in the suppressor strain.

**Importance:** There is increasing evidence that protein electron carriers like Fd have evolved to form specific partnerships with select electron donors and acceptors to keep native electron transfer pathways insulated from one another. This makes it challenging to integrate a Fd-dependent pathway like biological nitrogen fixation into non-nitrogen-fixing organisms and provide the high-energy reducing power needed to fix nitrogen. Here we show that amino acid substitutions in an electron donor for anaerobic aromatic compound degradation and a Fd involved in nitrogen fixation enabled electron transfer to nitrogenase. This work provides a model system to understand electron transfer chain specificity and how new electron transfer pathways can be evolved for biotechnologically valuable pathways like nitrogen fixation.

## Introduction

Ferredoxins (Fds) and flavodoxins (Flds) are small protein electron carriers that transfer a single electron from an electron donor to an electron acceptor. In particular, Fds have specialized over evolutionary time to associate with specific partner proteins, allowing Fds to selectively shuttle electrons to specific pathways (1, 2). Key factors such as the structure and charge of the Fd binding surface (3), regulation of Fd abundance (4), and the reduction potential of the Fd (5–7) all affect which partner proteins interact with a Fd. In theory, these properties can be altered to enable a Fd to interact with a new partner protein(s) to re-route electron flow, but this remains a challenge for rational design since these properties are ill-defined for many Fds. However, Fds can mediate electron transfer in biological reactions essential for growth, making it possible to select mutations that would allow a Fd and a new partner protein to interact (2, 8).

One such reaction is biological nitrogen fixation, which is catalyzed by the enzyme nitrogenase. Nitrogenase reduces atmospheric dinitrogen into ammonia and hydrogen using large amounts of ATP and low potential electrons delivered by Fd or Fld (9). In the purple non-sulfur bacterium, *Rhodopseudomonas palustris*, electron transfer to nitrogenase requires the FixABCX complex, which couples the oxidation of NADH to the reduction of quinone and a Fd or Fld using flavin-based electron bifurcation (Fig. 1) (10–12). *R. palustris* with a deletion in *fixA, fixB, fixC*, or *fixX* has a severe growth defect under nitrogen-fixing conditions, despite *R. palustris* encoding multiple Fd-reducing enzymes including pyruvate:Fd oxidoreductase (PFOR) and Fd-NAD(P)^+^ reductase (FNR) that play a role in electron transfer to nitrogenase in other diazotrophs (13–20). *R. palustris* also encodes six 2[4Fe-4S] Fds, with the primary electron donor to nitrogenase being the Fd, Fer1 (Rpa4631), although the Fld, FldA (Rpa2117), can also act as an electron donor in the absence of Fer1 and plays a role under iron-limiting conditions (11). Since *R. palustris* encodes multiple Fds and Fd-reducing enzymes, *R. palustris* in which FixABCX is inactive could be leveraged to select for mutations that would enable a new electron transfer chain (ETC) to nitrogenase.

**Figure 1.**
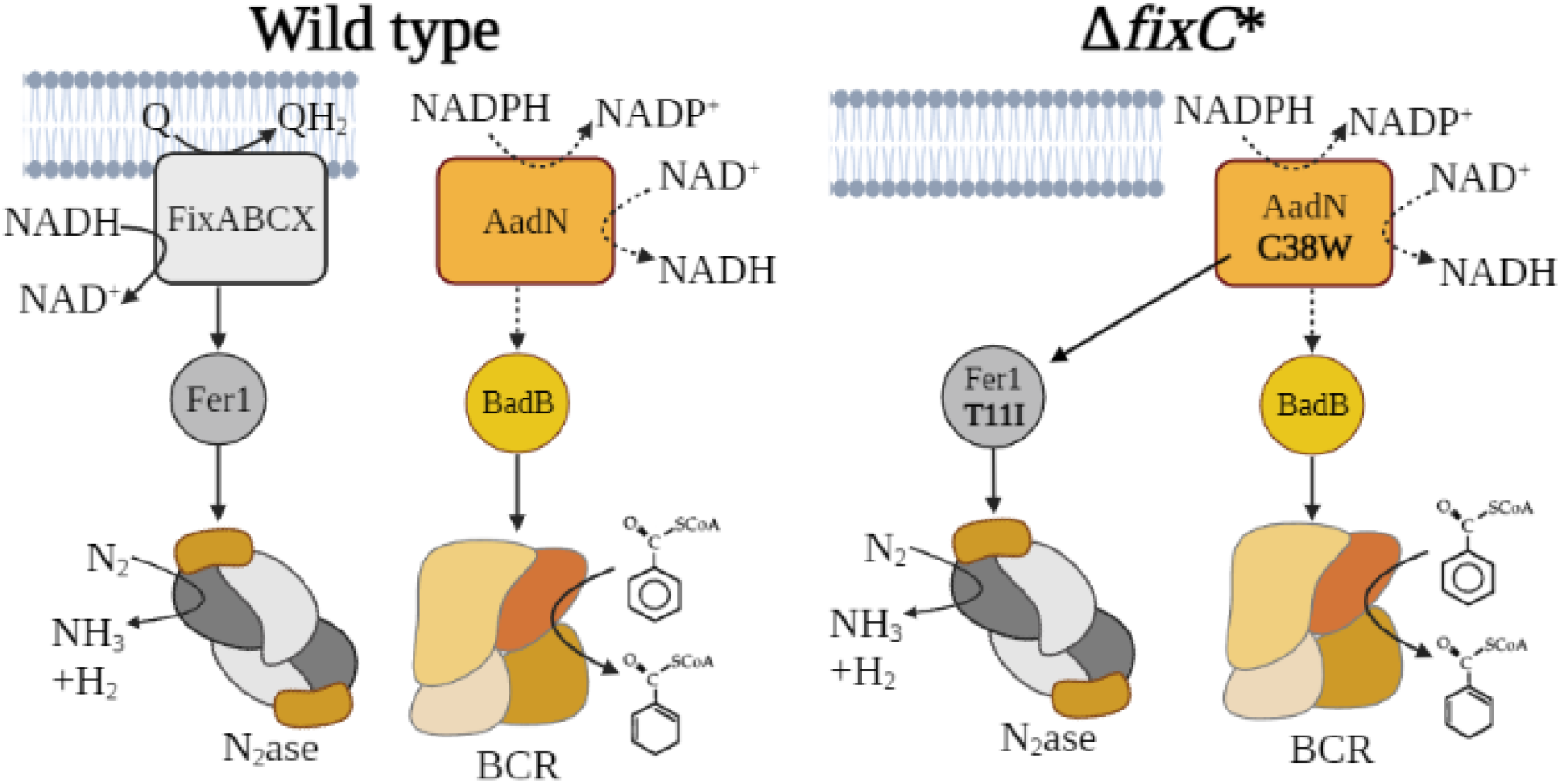
Current model of electron transfer to nitrogenase (N_2_ase) and benzoyl-CoA reductase (BCR) in *R. palustris*. Electron transfer to BCR is incompatible with nitrogen fixation in the wild type, but the C38W substitution in AadN and T11I substitution in Fer1 enable the electron transfer pathway for benzoate degradation to support nitrogen fixation in Δ*fixC\**. The hypothesized activity of AadN shown in dotted lines is inferred based on its similarity to NAD^+^- dependent NADPH:ferredoxin oxidoreductase from *Pyrococcus furiosis*.

Using an *R. palustris* Δ*fixC* strain, we isolated a suppressor mutant that restored growth of this strain under nitrogen-fixing conditions. We found two mutations in the suppressor strain were required to restore nitrogenase activity in the absence of a functional FixABCX complex (Fig. 1). One mutation was in the gene encoding Fer1, while the second was in an uncharacterized gene *rpa0678*. Sequence and genetic analysis revealed that the protein encoded by this gene is a homolog of a flavin-based electron bifurcating NAD^+^-dependent Fd:NADPH oxidoreductase (Nfn), and we found it is required for anaerobic aromatic compound degradation in *R. palustris* (Fig. 1). Because of its role in anaerobic aromatic compound degradation, we have renamed *rpa0678* to *aadN* for anaerobic aromatic degradation, Nfn-like protein. The data herein support a model where a new ETC for nitrogenase formed between components of two endogenous ETCs and provides a system that can be used to study the determinants of selective electron transfer.

## Results

### A mutation in fer1 is necessary but not sufficient to support electron transfer to nitrogenase in the absence of FixC

As shown in Fig. 2A, *R. palustris* Δ*fixC* has a severe growth defect when grown under nitrogen-fixing conditions. To select for suppressor mutants of *R. palustris ΔfixC*, this strain was incubated in nitrogen-fixing conditions for several weeks in the light. One of three replicate liquid cultures grew, from which a suppressor mutant of *R. palustris* Δ*fixC* was isolated, referred hereafter as *R. palustris* Δ*fixC\**. Deletion of *fixA* in *R. palustris* Δ*fixC\** did not disrupt the ability of the suppressor strain to grow in nitrogen-fixing conditions, confirming that the remaining Fix complex is not required in *R. palustris* Δ*fixC\** (Fig. 2A). Genome sequencing revealed that Δ*fixC\** accumulated 18 mutations in 16 different genes (Table 1). One of the mutations was in *recQ*, a DNA helicase involved in DNA repair, which may account for the large number of mutations found in the suppressor strain (21). While most of the mutations did not have an obvious connection to electron transfer, one of the mutations identified was in *fer1*, which encodes the primary electron donor to nitrogenase in *R. palustris*.

**Figure 2.**
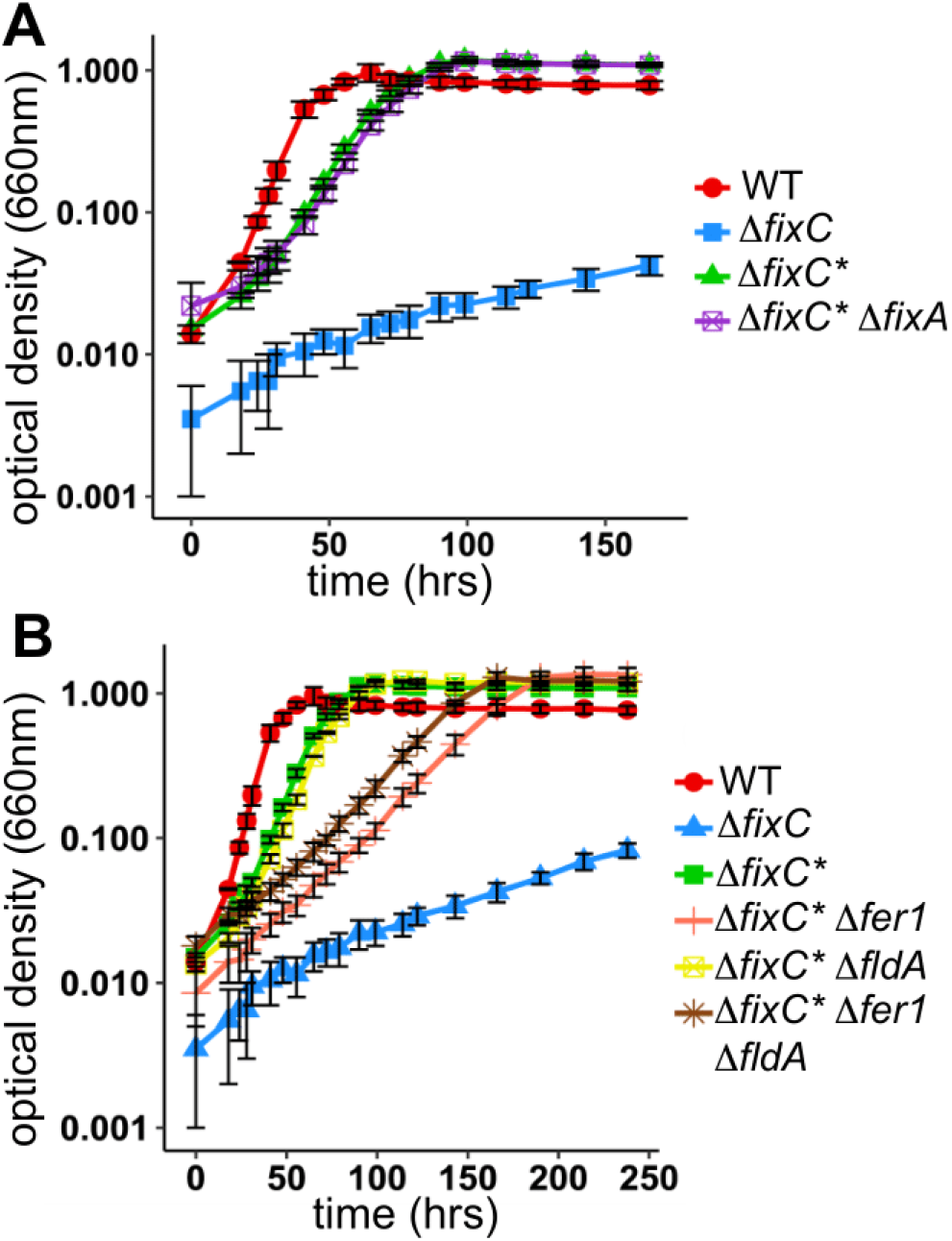
Growth of Δ*fixC\** in nitrogen-fixing conditions does not require *fixA* but does require *fer1^T11I^*. (**A**) Growth of wild-type *R. palustris* (WT), *R. palustris* with a deletion in *fixC* (Δ*fixC*), *R. palustris* suppressor of Δ*fixC* (Δ*fixC\**), and *R. palustris* suppressor of Δ*fixC* with a deletion in *fixA* (Δ*fixC\** Δ*fixA*) in minimal medium lacking ammonium sulfate (nitrogen-fixing) with 20 mM acetate. (**B**) Growth of wild-type *R. palustris* (WT), *R. palustris* with a deletion in *fixC* (Δ*fixC*), *R. palustris* suppressor of Δ*fixC* (Δ*fixC\**), *R. palustris* suppressor of Δ*fixC* with a deletion in *fer1* (Δ*fixC\** Δ*fer1*), *R. palustris* suppressor of Δ*fixC* with a deletion in *fldA* (Δ*fixC\** Δ*fldA*), and *R. palustris* suppressor of Δ*fixC* with a deletion in *fer1* and *fldA* (Δ*fixC\** Δ*fer1* Δ*fldA*) in minimal medium lacking ammonium sulfate (nitrogen-fixing) with 20 mM acetate. For both **A** and **B** data are the average of two biological replicates, error bars represent one standard deviation from the mean.

**Table 1.**
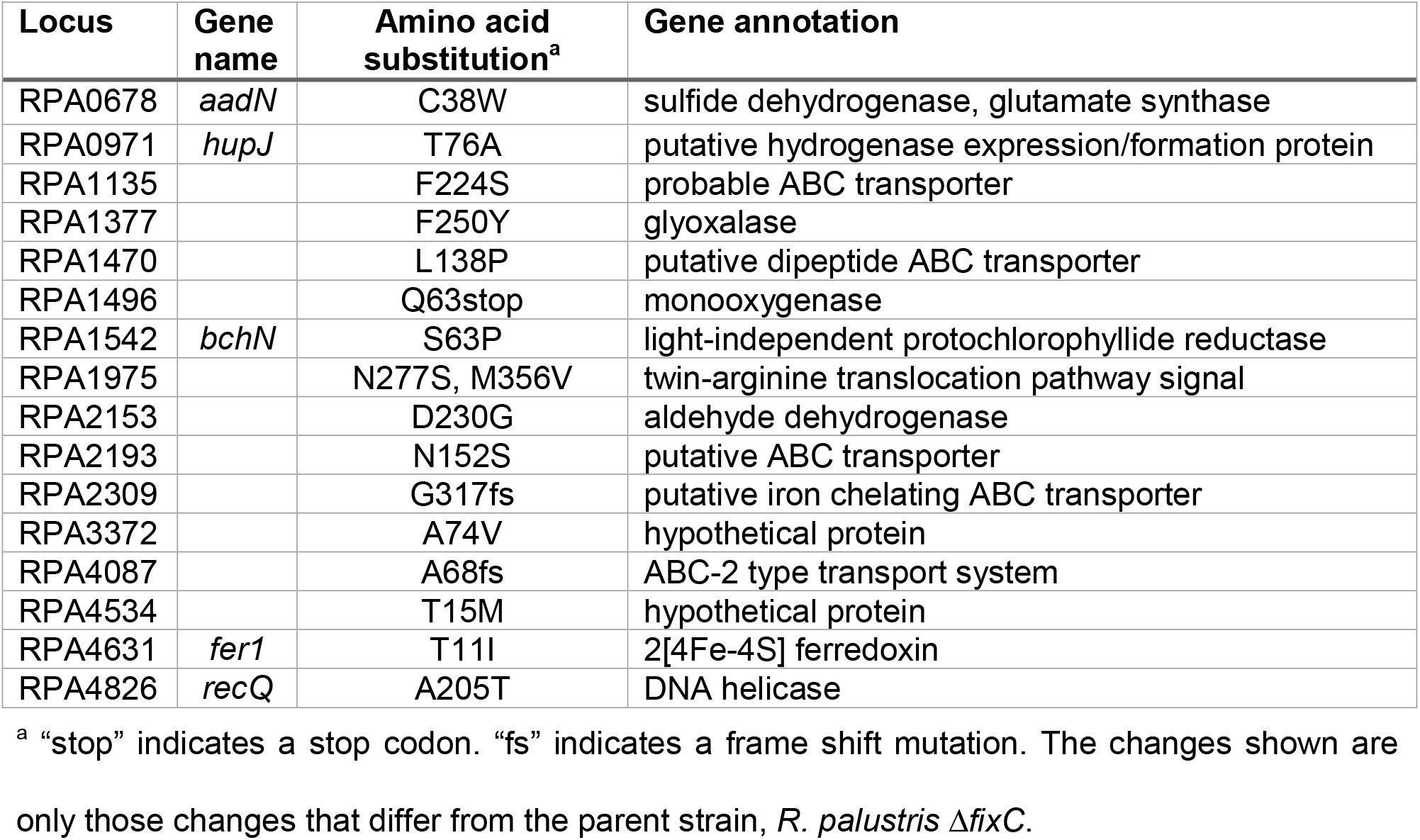
Mutations found in the genome of Δ*fixC\** by whole genome sequencing.

| Locus | Gene name | Amino acid substitution<sup>a</sup> | Gene annotation |
| --- | --- | --- | --- |
| RPA0678 | *aadN* | C38W | sulfide dehydrogenase, glutamate synthase |
| RPA0971 | *hupJ* | T76A | putative hydrogenase expression/formation protein |
| RPA1135 |  | F224S | probable ABC transporter |
| RPA1377 |  | F250Y | glyoxalase |
| RPA1470 |  | L138P | putative dipeptide ABC transporter |
| RPA1496 |  | Q63stop | monooxygenase |
| RPA1542 | *bchN* | S63P | light-independent protochlorophyllide reductase |
| RPA1975 |  | N277S, M356V | twin-arginine translocation pathway signal |
| RPA2153 |  | D230G | aldehyde dehydrogenase |
| RPA2193 |  | N152S | putative ABC transporter |
| RPA2309 |  | G317fs | putative iron chelating ABC transporter |
| RPA3372 |  | A74V | hypothetical protein |
| RPA4087 |  | A68fs | ABC-2 type transport system |
| RPA4534 |  | T15M | hypothetical protein |
| RPA4631 | *fer1* | T11I | 2[4Fe-4S] ferredoxin |
| RPA4826 | *recQ* | A205T | DNA helicase |

<sup>a</sup> “stop” indicates a stop codon. “fs” indicates a frame shift mutation. The changes shown are only those changes that differ from the parent strain, *R. palustris* $\Delta fixC$ .

The mutation in *fer1* results in the substitution of threonine 11 for isoleucine (T11I). To determine if the Fer1 T11I variant was required for the suppressor phenotype, the *fer1^T11I^* allele in *R. palustris* Δ*fixC\** was replaced with either an in-frame deletion in *fer1* or a wild-type *fer1* allele. Both strains had a significantly slower growth rate than *R. palustris* Δ*fixC\** (Table 2 and Fig. S1), indicating that the *fer1^T11I^* allele is required for the suppressor phenotype. However, even in the absence of Fer1, the suppressor strain was still able to grow, albeit at a reduced rate, suggesting that other electron carriers can compensate in the absence of Fer1.

**Table 2.**
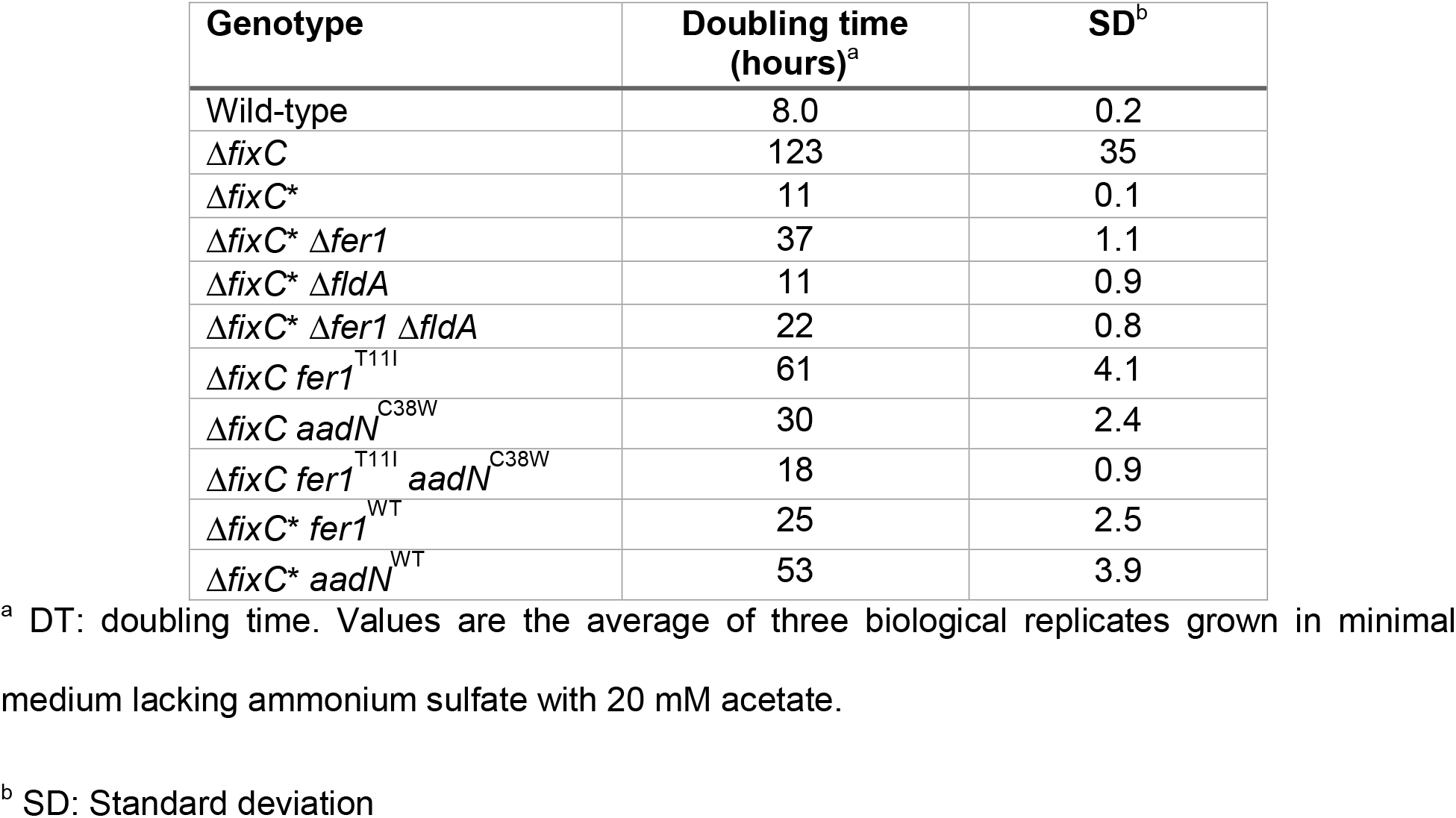
Growth rates of *R. palustris* strains in nitrogen-fixing conditions.

| Genotype | Doubling time (hours)<sup>a</sup> | SD<sup>b</sup> |
| --- | --- | --- |
| Wild-type | 8.0 | 0.2 |
| $\Delta fixC$ | 123 | 35 |
| $\Delta fixC^*$ | 11 | 0.1 |
| $\Delta fixC^* \Delta fer1$ | 37 | 1.1 |
| $\Delta fixC^* \Delta fldA$ | 11 | 0.9 |
| $\Delta fixC^* \Delta fer1 \Delta fldA$ | 22 | 0.8 |
| $\Delta fixC fer1^{T11I}$ | 61 | 4.1 |
| $\Delta fixC aadN^{C38W}$ | 30 | 2.4 |
| $\Delta fixC fer1^{T11I} aadN^{C38W}$ | 18 | 0.9 |
| $\Delta fixC^* fer1^{WT}$ | 25 | 2.5 |
| $\Delta fixC^* aadN^{WT}$ | 53 | 3.9 |

<sup>a</sup> DT: doubling time. Values are the average of three biological replicates grown in minimal medium lacking ammonium sulfate with 20 mM acetate.
<sup>b</sup> SD: Standard deviation

To determine if the *fer1^T11I^* allele was sufficient to restore growth under nitrogen-fixing conditions in the absence of an intact FixABCX complex, the *fer1^T11I^* allele was introduced into *R. palustris ΔfixC*. As shown in Fig. 3, the *fer1^T11I^* allele did not restore growth in *R. palustris* Δ*fixC* under nitrogen-fixing conditions, indicating other mutations in *R. palustris* Δ*fixC\** are required for electron transfer. This finding revealed that the *fer1*^T11I^ allele was necessary but not sufficient for the suppressor phenotype. Because many of the remaining mutations in *R. palustris* Δ*fixC\** had unknown or hypothetical functions (Table 1) we needed to broaden our search for genes involved in electron transfer to nitrogenase in *R. palustris* Δ*fixC\**.

**Figure 3:**
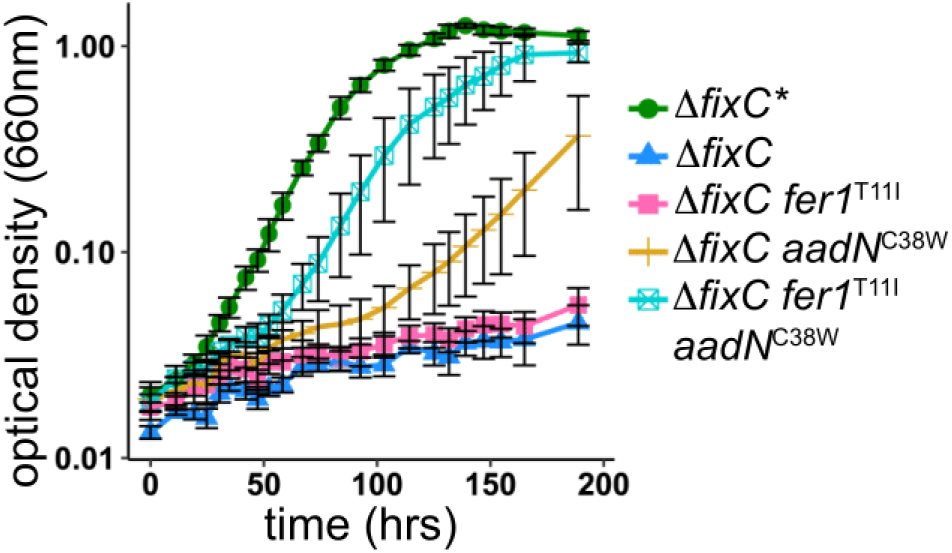
Δ*fixC\** alleles of *fer1* and *aadN* enable growth of Δ*fixC* in nitrogen fixing conditions. Growth of *R. palustris* suppressor of Δ*fixC* (Δ*fixC\**), *R. palustris* with a deletion in *fixC* (Δ*fixC*), *R. palustris* Δ*fixC* encoding a *fer7*^T11I^ allele (Δ*fixC fer1*^T11I^), *R. palustris* Δ*fixC* encoding an *aadN*^C38W^ allele (Δ*fixC aadN*^C38W^), and *R. palustris* Δ*fixC* encoding the *fer1^T11I^* and *aadN*^C38W^ alleles (Δ*fixC fer1*^T11I^ *aadN*^C38W^) in minimal medium lacking ammonium sulfate (nitrogen-fixing) supplemented with 20 mM acetate. Data shown are the average of three biological replicates, error bars represent one standard deviation from the mean.

Nitrogenase is a very energetically expensive reaction, requiring at least 8 low-potential electrons per catalytic cycle (9, 22). We hypothesized that any components of the ETC would be transcribed at higher rates to help accommodate the demand for reducing power in *R. palustris* Δ*fixC\**. Therefore, RNA-seq analysis was carried out to compare gene expression changes in *R. palustris* Δ*fixC\** compared to wild-type *R. palustris* under nitrogen-fixing conditions. We found that expression of *fldA* had the highest change in gene expression and was up-regulated 23-fold in *R. palustris* Δ*fixC\** compared to wild-type *R. palustris* (supplemental dataset 1). To test the role of FldA in the ETC used by *R. palustris* Δ*fixC\**, strains were constructed with in-frame deletions in *fldA* and their growth rates in nitrogen-fixing conditions were measured (Fig. 2B, Table 2). We found FldA was not required for *R. palustris* Δ*fixC\** to grow under nitrogen-fixing conditions, and FldA was not redundant with Fer1 in *R. palustris* Δ*fixC\** since there was no significant difference in growth rate between *R. palustris* Δ*fixC\** strains with a deletion in *fer1* or both *fer1* and *fldA* (Fig. 2B). RNA-seq also showed that most other genes encoding enzymes known to reduce Fd were down-regulated or showed relatively minor (less than 2-fold) changes in gene expression in *R. palustris* Δ*fixC\** (Table S1).

### A mutation in aadN is both necessary and sufficient for electron transfer to nitrogenase in the absence of FixC

To identify which genes are required for electron transfer to nitrogenase in *R. palustris* Δ*fixC\**, we used a random transposon mutagenesis strategy combined with a metronidazole enrichment (Fig. S2) (23). Metronidazole is an antibiotic that is activated when reduced by low-potential electron carriers, specifically Fd and Fld, causing cell death (24). Transposon mutants that survive metronidazole enrichment likely have insertions that disrupt electron transfer. Using this approach, we identified one transposon mutant that survived the enrichment and grew similar to *R. palustris* Δ*fixC\** under non-nitrogen-fixing conditions but could not grow under nitrogen-fixing conditions. This mutant had a transposon insertion in *aadN (rpa0678*). In *R. palustris ΔfixC*, aadN* had a non-synonymous mutation, encoding a variant of AadN in which cysteine 38 is substituted for tryptophan (C38W) (Table 1). We found that replacing the *aadN*^C38W^ allele in *R. palustris* Δ*fixC\** with wild-type *aadN* disrupted the ability of the strain to grow in nitrogen-fixing conditions (Table 2). When the *aadN*^C38W^ mutation was introduced into the parent strain, *R. palustris ΔfixC*, this mutation alone allowed *R. palustris* Δ*fixC* to grow under nitrogen-fixing conditions, indicating that the *aadN*^C38W^ mutation is both necessary and sufficient to restore growth of *R. palustris* Δ*fixC* under nitrogen-fixing conditions. However, the growth rate of *R. palustris ΔfixC aadN*^C38W^ was slower than *R. palustris* Δ*fixC\** (Fig. 3 and Table 2). When the *aadN*^C38W^ mutation was combined with the *fer1*^T11I^ mutation in *R. palustris ΔfixC*, the growth rate increased (Fig. 3 and Table 2). This suggests that the variants of AadN and Fer1 form a new ETC that can deliver electrons to nitrogenase in the absence of FixABCX (Fig. 1).

We measured hydrogen production in growing cultures to quantify nitrogenase activity. Hydrogen is an obligate product of nitrogenase activity (9). Nitrogenase activity can be estimated by measuring hydrogen production in these strains because they have a defect in the expression of an uptake hydrogenase and accumulate hydrogen under nitrogen-fixing conditions (25). Introduction of the *aadN*^C38W^ allele into *R. palustris* Δ*fixC* was sufficient to allow hydrogen production, but it produced about 50 percent less hydrogen compared to wild-type *R. palustris* or *R. palustris* Δ*fixC\** (Table 3). However, when both the *fer1*^T11I^ and *aadN*^C38W^ alleles were introduced into *R. palustris ΔfixC*, hydrogen production was restored to levels observed for *R. palustris* Δ*fixC\**, confirming that only these two mutations are required for electron transfer to nitrogenase in the absence of a functional FixABCX complex (Table 3).

**Table 3.**
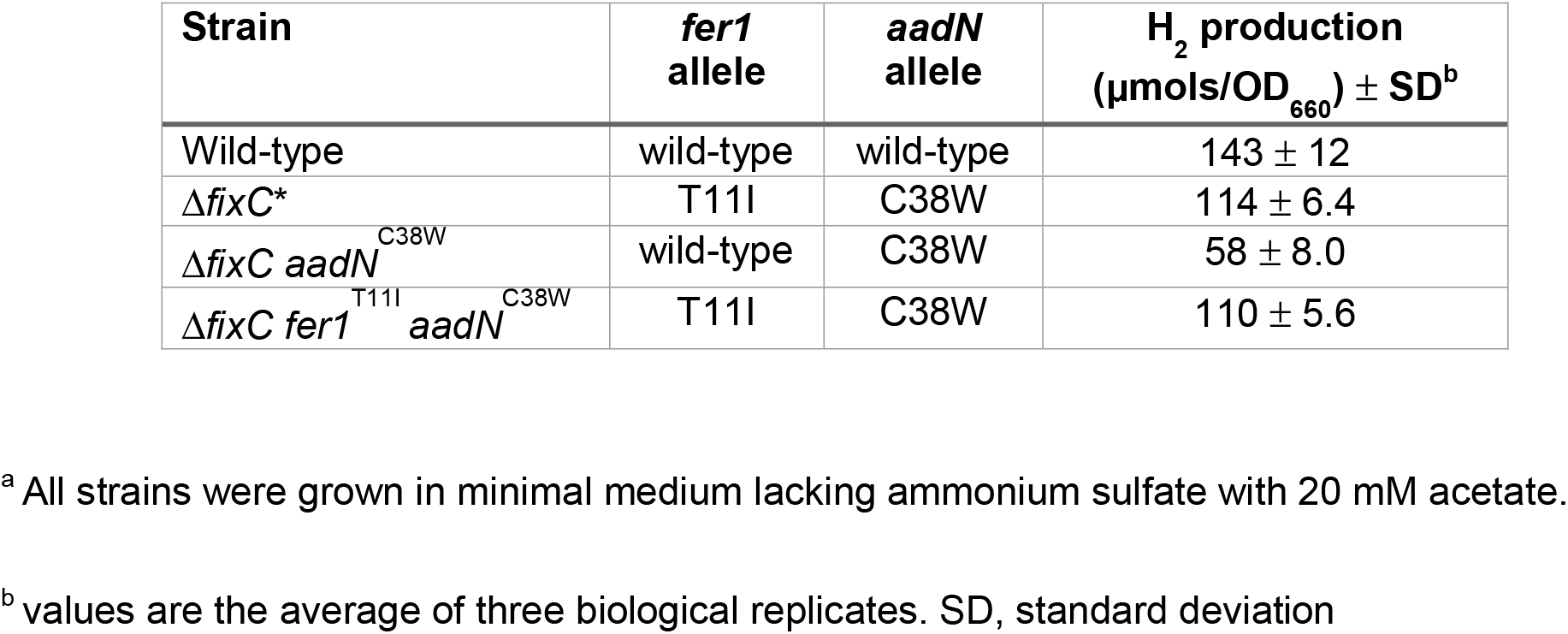
Hydrogen production of *R. palustris* strains grown under nitrogen-fixing conditions^a^.

| Strain | *fer1*<br>allele | *aadN*<br>allele | H<sub>2</sub> production<br>(μmols/OD<sub>660</sub>) ± SD<sup>b</sup> |
| --- | --- | --- | --- |
| Wild-type | wild-type | wild-type | 143 ± 12 |
| Δ*fixC*<sup>*</sup> | T11I | C38W | 114 ± 6.4 |
| Δ*fixC* *aadN*<sup>C38W</sup> | wild-type | C38W | 58 ± 8.0 |
| Δ*fixC* *fer1*<sup>T11I</sup> *aadN*<sup>C38W</sup> | T11I | C38W | 110 ± 5.6 |

<sup>a</sup> All strains were grown in minimal medium lacking ammonium sulfate with 20 mM acetate.
<sup>b</sup> values are the average of three biological replicates. SD, standard deviation

### AadN is a homolog of a Fd-reducing enzyme and is required for anaerobic aromatic compound degradation

While these data indicate that AadN^C38W^ is required for electron transfer to nitrogenase in *R. palustris* Δ*fixC\**, the native function of AadN in *R. palustris* was unclear. To gain insight into how AadN may play a role in electron transfer, we analyzed the protein domain organization and used protein modeling to make predictions about the structure and activity of AadN (26, 27). Although AadN is annotated as a sulfide dehydrogenase, sequence analysis of AadN revealed that it shares homology with both the large and small subunit of the enzyme NfnI from *Pyrococcus furiosis* (*Pf*NfnI, Fig. 4A) (28). *Pf*NfnI is an NAD^+^-dependent Fd:NADPH oxidoreductase (Nfn) that uses flavin-based electron bifurcation to balance NADP(H), NAD(H) and Fd pools to conserve energy and maintain redox balance (Fig. 4B) (28, 29). While the large and the small subunit of *Pf*NfnI are encoded by two separate genes, these subunits are fused in AadN and the domain organization of AadN is more like concatenated, pattern B Nfn as described in (30) (Fig. 4A). We found the cofactor binding domains in *Pf*NfnI are conserved in AadN and both the large and small subunits share 51% or more amino acid identity (Fig 4A). This suggests that AadN may be able to carry out flavin-based electron bifurcation and use electrons from NAD(P)H to reduce Fd (Fig. 4B).

**Figure 4.**
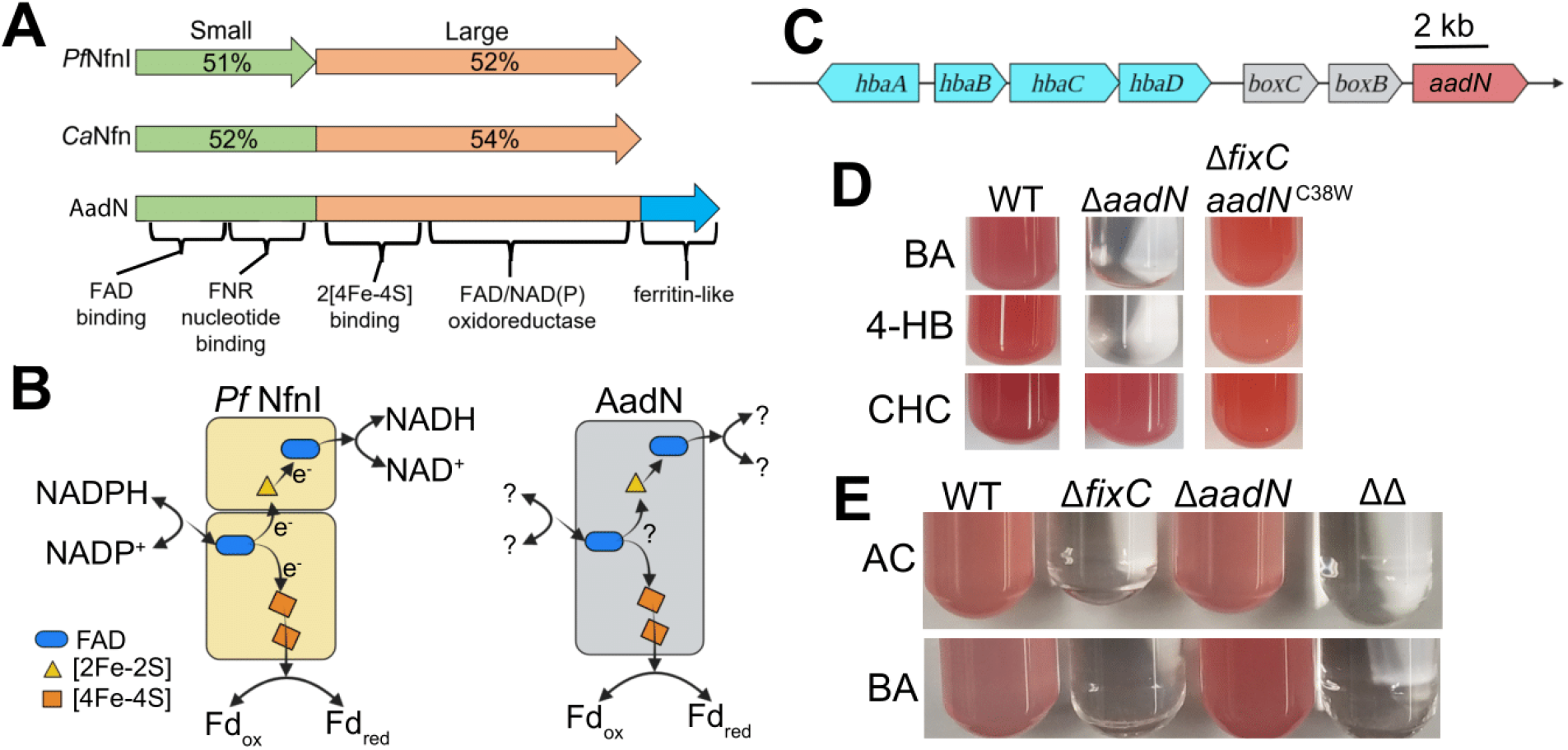
AadN, an Nfn homolog, is required for anaerobic aromatic compound degradation. (**A**) AadN is homologous to NfnI from *P. furiosis* (*Pf*NfnI) and a Pattern B Nfn from *Clostridium autoethanogenum* (*Ca*Nfn). The percent amino acid identity of the large and small subunits of *Pf*NfnI and *Ca*Nfn to AadN are shown over the small (green) and large (orange) regions. (**B**) *Pf*NfnI ligates two [4Fe-4S] clusters, one [2Fe-2S] cluster, and two flavin adenine dinucleotide (FAD) cofactors per *Pf*NfnI heterodimer. The amino acid sequence of the binding domains for each of these cofactors is conserved in AadN, but it remains unclear if AadN interacts with the same redox pools as *Pf*NfnI. (**C**). Map of genomic region around *aadN* in *R. palustris* CGA009 shows that *aadN* is near genes involved in aromatic compound degradation (*hba* genes, cyan). (**D**) Growth phenotypes of *R. palustris* (WT), *R. palustris* with a deletion in *aadN (ΔaadN*), and *R. palustris* with a deletion in *fixC* encoding the *aadN*^C38W^ allele (Δ*fixC 0678*^C38W^) in minimal medium supplemented with 10 mM HCO_3_ and either 5.7 mM benzoate (BA), 5.7 mM 4-hydroxybenzoate (4-HB), or 5.7 mM cyclohexane carboxylate (CHC) as carbon sources. Cultures shown are representative of three independent trials. (**E**) Growth phenotypes of *R. palustris* (WT), *R. palustris* with a deletion in *fixC (ΔfixC), R. palustris* with a deletion in *aadN (ΔaadN*) and *R. palustris* with a deletion in both *fixC* and *aadN* (ΔΔ) in minimal medium lacking ammonium sulfate (nitrogen-fixing) supplemented with 20 mM acetate (AC) or 5.7 mM benzoate (BA). Cultures shown are representative of three independent trials.

We also found that *aadN* is adjacent to genes involved in anaerobic degradation of aromatic compounds such as benzoate and 4-hydroxybenzoate (4-HB) (Fig. 4C) (31). Anaerobic degradation of these compounds requires the enzyme benzoyl-CoA reductase, which carries out ATP-dependent electron transfer from a low potential Fd to reduce a CoA thioester in benzoyl-CoA (Fig. S3) (32–34). In *Thauera aromatica*, benzoyl-CoA reductase is supplied with reducing power through a Fd:2-oxoglutarate oxidoreductase known as KorAB (35). While two strains of *R. palustris* encode *korAB* homologs in the same gene cluster as other benzoate degradation genes, seven *R. palustris* strains that encode genes for anaerobic aromatic compound degradation lack *korAB* (36). If AadN plays a role in electron transfer to benzoyl-CoA reductase, we reasoned that strains lacking *korAB* should encode *aadN*. Among *R. palustris* strains, *aadN* is present in the genomes of the six strains lacking *korAB* but is not found in the two strains encoding *korAB* (Table 4). This suggests that *R. palustris* strains either use AadN or KorAB for electron transfer during aromatic compound degradation but not both (Table 4). The only *R. palustris* strain lacking *aadN* and *korAB* was strain HaA2, which cannot degrade aromatic compounds (36).

**Table 4.** Genetic distribution of *aadN* and *korAB* among *R. palustris* strains.

| *R. palustris*<br>strain | *korAB*<sup>a</sup> | *aadN*<sup>b</sup> |
| --- | --- | --- |
| CGA009 | - | + |
| TIE-1 | - | + |
| DX-1 | - | + |
| DCP3 | - | + |
| BisB18 | - | + |
| PS3 | - | + |
| YSC3 | - | + |
| BisB5 | + | - |
| BisA53 | + | - |
| HaA2<sup>c</sup> | - | - |

<sup>a</sup> +, contains a homolog of *korA* (WP\_011501953.1) from *R. palustris* BisB5 with greater than 80% amino acid identity within the anaerobic aromatic degradation gene cluster; -, no homolog present.
<sup>b</sup> +, contains a homolog of *aadN* (WP\_011156245) with greater than 90% amino acid identity within the anaerobic aromatic degradation gene cluster; -, no homolog present.
<sup>c</sup> *R. palustris* strain HaA2 is unable to degrade aromatic compounds.

Given its proximity to genes required for anaerobic aromatic compound degradation, its similarity to *Pf*NfnI, and its importance in electron transfer to nitrogenase in Δ*fixC\**, we hypothesized that AadN plays a role in electron transfer to benzoyl-CoA reductase. To test this, we looked at the ability of *R. palustris ΔaadN* to metabolize aromatic compounds. Benzoate and 4-HB are converted to benzoyl-CoA and are reductively dearomatized by benzoyl-CoA reductase, but cyclohexane carboxylate (CHC) enters the same degradation pathway after this de-aromatization step (Fig. S3) (36). When benzoate or 4-HB were provided as sole carbon sources, *R. palustris ΔaadN* had a growth defect, indicating that *aadN* is required for degradation of benzoate and 4-HB (Fig. 4D). We found that AadN is not required to grow on CHC because both wild-type *R. palustris* and *R. palustris ΔaadN* were able to grow when CHC was provided as the sole carbon source (Fig. 4D). We also found that the C38W substitution in AadN did not disrupt the ability of *R. palustris aadN*^C38W^ to grow on aromatic carbon sources, indicating that the C38W variant is able to play a role in both nitrogen fixation and anaerobic aromatic compound degradation (Fig. 4D). In tandem with evidence from protein sequence analysis, this suggests that AadN plays a role in electron transfer to benzoyl-CoA reductase.

### The native ETC for anaerobic aromatic compound degradation is insulated from nitrogen fixation

Since *R. palustris* Δ*fixC* is unable to grow under nitrogen-fixing conditions, the ETC for anaerobic aromatic compound degradation and nitrogen fixation are likely insulated from each other. To probe the insulation of these two ETCs, we grew *R. palustris* strains with an in-frame deletion in *fixC* or *aadN* under nitrogen-fixing conditions with benzoate as a sole carbon source. As shown in Fig. 4E, we found that although *aadN* was required for growth on benzoate under non-nitrogen-fixing conditions, *R. palustris ΔaadN* was able to grow with benzoate as a carbon source under nitrogen-fixing conditions, suggesting that FixABCX may be able to complement the loss of *aadN* and support benzoate degradation under nitrogen-fixing conditions (Fig. 4E). However, *R. palustris* Δ*fixC* was unable to grow under nitrogen-fixing conditions with benzoate, indicating that the ETC for benzoate degradation cannot sustain electron transfer to nitrogenase. This data supports a model in which electron transfer for benzoate degradation is insulated from nitrogen fixation, and the C38W substitution in AadN overcomes the apparent insulation to allow AadN to function in electron transfer to nitrogenase.

## Discussion

In this study, we wanted to identify mutations that would restore electron transfer to nitrogenase in *R. palustris* in the absence of its native electron donor, FixABCX. We hypothesized that changes in Fd or Fld would be required to alter the flow of electrons in the cell, enabling the formation of a new ETC from existing components. While we found that a single amino acid substitution in Fer1 was important for the new ETC, it was not sufficient on its own to support electron transfer to nitrogenase in the absence of FixABCX. We found that a mutation in an Nfn-like gene we termed *aadN* was sufficient for electron transfer, but when combined with the mutation in *fer1*, electron transfer to nitrogenase was more efficient. While changing the properties of a Fd played an important role in making the new ETC more efficient, a single change in a Fd-reducing enzyme had a larger role in the formation of the new ETC. Therefore, it is likely that changes in both the Fd-reducing enzyme and the Fd will be required to optimize electron transfer through engineered pathways.

Our approach also uncovered a new role for an uncharacterized Nfn homolog. Nfn homologs are found in all domains of life, but the physiological role for many of these homologs is unknown (30, 37). Sequence homology revealed that AadN is related to Nfn and is part of an entirely uncharacterized family of Nfn enzymes known as pattern B Nfns, in which the two subunits of Nfn are fused (30). Pattern A Nfns, including *Pf*NfnI, ligate two [4Fe-4S] clusters, one [2Fe-2S] cluster, two FAD cofactors, and have binding sites for NADPH and NAD^+^ (28, 38). We found that each of these substrate and cofactor binding sites were conserved in AadN, suggesting that AadN carries out FBEB using NADPH and NAD^+^ to reduce Fd. However, not all Nfn homologs show Nfn bifurcating activity (39), and further structural and enzymatic analysis will be required to determine if AadN carries out FBEB using the same substrates as Nfn. We showed that AadN is required for aromatic compound degradation and likely plays a role in electron transfer to benzoyl-CoA reductase. To the best of our knowledge, this is the first proposed role for a pattern B Nfn, and this discovery provides evidence that an Nfn-like enzyme can supply reducing power for anaerobic aromatic compound degradation. Our results also implicate the Nfn enzyme family in electron transfer to nitrogenase. Some diazotrophs do not appear to encode any Fd- or Fld-reducing enzymes known to be involved in electron transfer to nitrogenase (40). Our evidence that an Nfn homolog can supply reducing power to nitrogenase may help illuminate the ETCs for nitrogen fixation in some of these diazotrophic organisms.

The insulation of nitrogen fixation and anaerobic aromatic compound degradation highlights the complicated nature of electron transfer insulation. The key enzyme in anaerobic aromatic compound degradation, benzoyl-CoA reductase, requires low potential electrons delivered by a Fd (41). The benzoate degradation gene cluster in *R. palustris* encodes a Fd known as BadB, and a homolog of BadB in *T. aromatica* has been shown to have a very low midpoint potential of −587 mV (31, 34). Based on thermodynamics alone, the ETC for anaerobic aromatic compound degradation is predicted to be compatible with nitrogenase. However, we found that even when grown with benzoate, *R. palustris* Δ*fixC* could not grow under nitrogen-fixing conditions, indicating that the ETC for benzoyl-CoA reductase cannot support electron transfer to nitrogenase.

The electron transfer insulation we observe could be due, in part, to the inability of BadB to interact with nitrogenase. However, our results indicate that the insulation between these two pathways must also be due to the inability of AadN to reduce Fer1 since an amino acid substitution in AadN restores some activity to nitrogenase and is further facilitated by an amino acid substitution in Fer1. It is unclear how these variants enable electron transfer to nitrogenase, but it is likely that they facilitate interaction between AadN and Fer1. The C38W amino acid substitution in AadN could enable electron transfer to nitrogenase by disrupting post-translational regulation of AadN, affecting the stability of AadN, or altering the Fd binding site of AadN. The threonine residue at position 11 in Fer1 is adjacent to a cysteine predicted to coordinate one of the [4Fe-4S] clusters in Fer1. Many other low potential 2[4Fe-4S] Fds encode isoleucine at this position and similar threonine to isoleucine substitutions in other Fds have been shown to lower the reduction potential of Fds (5, 34, 42). This suggests that the T11I substitution in Fer1 lowers its reduction potential, although it is unclear why this would facilitate interaction with AadN. Further characterization of how these amino acid substitutions alter the properties of these proteins is needed to understand how these proteins can form a new electron transfer pathway to nitrogenase.

In summary, this study illustrates the potential of using a selection strategy to enable a new electron transfer pathway to nitrogenase. Given that electron transfer to nitrogenase is a major hurdle to engineering non-nitrogen-fixing organisms to fix nitrogen, this approach could be useful in evolving non-native ETCs to be compatible with nitrogenase.

## Materials & Methods

### Reagents, bacteria, and culture methods

All *R. palustris* strains were grown in a defined mineral medium (non-nitrogen-fixing medium) or mineral medium lacking ammonium sulfate (nitrogen-fixing medium) as described previously (43). Media were prepared using an anaerobic chamber (atmosphere: 98% N_2_, 2% H_2_, <10 ppm O_2_) as described previously (11). Liquid cultures were supplemented with 20 mM acetate and agar plates were supplemented with 10 mM succinate as carbon sources. Where indicated, liquid cultures were grown with 5.7 mM benzoate, 4-hydroxybenzoate, or cyclohexane carboxylate and were supplemented with 10 mM HCO_3_ (44). Plates were incubated in GasPak™ EZ anaerobe container systems at 30 °C (Becton Dickinson). Plates were placed within 10 inches of a 60W incandescent light bulb and liquid cultures were placed within 5.5 inches of the light bulb, which provides 30 μmol photons m^-2^s^-1^ (General Electric). Where applicable, *R. palustris* was grown with 100 μg/mL gentamicin and 200 μg/mL kanamycin, and *E. coli* strains were grown in lysogeny broth (LB) at 37°C supplemented with gentamicin (20 μg/mL). For metronidazole enrichment, metronidazole was added to a final concentration of 50 mM.

### Genetic manipulation of R. palustris

For each gene of interest, a corresponding pJQ200SK-derived deletion or allelic exchange vector was created (Table S2) (45). Deletion vectors included approximately 1 kb of sequence upstream of the start codon and 1 kb of sequence downstream of the stop codon of the gene to be removed. Allelic exchange vectors contained 1 kb of sequence upstream and downstream of the point mutation of interest. The deletion or allelic exchange vectors were mobilized into *R. palustris* by conjugation using *E. coli* S17-1 using standard methods (46). Gene deletions were confirmed by PCR using primers designed to bind upstream and downstream of the gene of interest (Table S2). PCR products were analyzed using gel electrophoresis with a 1% agarose gel. All allelic exchange strains were confirmed by Sanger sequencing (GENEWIZ, South Plainfield, NJ).

### RNA extraction, cDNA library preparation, and sequencing

*R. palustris* cells were harvested from 10 mL of liquid nitrogen-fixing medium after cultures had grown to an optical density (660 nm) of 0.45-0.47. Cells were incubated on ice for 10 minutes then harvested by centrifugation. The cell pellet was flash frozen in liquid nitrogen and stored at −80°C. Cell pellets were thawed and resuspended in 1 mL QIAzol Lysis Reagent (QIAGEN, Hilden, Germany) and homogenized using a BioSpec Products BeadBeater-24 (Bartlesville, OK) at maximum rpm for one minute at 4°C, then allowed to cool for one minute on ice. This cycle was repeated four times to fully homogenize the sample. Total RNA was isolated using the miRNAeasy Mini Kit (QIAGEN, Hilden, Germany), and DNA was removed by incubating each sample with TURBO DNase (Invitrogen, Carlsbad, CA). Following DNA depletion, RNA was purified and concentrated using the RNeasy MiniElute Cleanup Kit (QIAGEN, Hilden, Germany). cDNA library construction and library sequencing were performed at GENEWIZ, LLC (South Plainfield, NJ). During cDNA library construction, ribosomal RNA was depleted using the Ribo-Zero rRNA removal kit (Illumina, San Diego, CA). cDNA was prepared using the NEBNext Ultra RNA Library Prep Kit and sequencing reactions, image analysis, and base calling were all performed on an Illumina HiSeq 2500 instrument (Illumina, San Diego, CA).

### Differential Gene Expression Analysis

Quality base calling in sequencing data was analyzed using the FastQC application (v 0.11.8, https://www.bioinformatics.babraham.ac.uk/projects/fastqc/) and TrimGalore (v 0.6.2) was used to remove adapter sequences, process, and validate all reads using the default parameters. Further analysis was performed on the Avadis software package (v 3.1.1, Strand Life Sciences, Bengaluru, India). Reads were aligned to the published genome of *R. palustris* CGA009 and differentially expressed genes were identified using the DESeq2 package (47) in R version 3.6 using the default parameters.

### Transposon mutagenesis and metronidazole enrichment

Cultures of *E. coli* BW20767 (48) and *R. palustris* Δ*fixC\** were grown to mid-log phase, washed with minimal medium twice, mixed at equivalent concentrations, then plated on minimal medium agar supplemented with 10 mM succinate, 0.2% yeast extract, and 0.5% casamino acids. Plates were incubated overnight at 30°C. Following incubation, all visible biomass was transferred from the plate into liquid minimal medium supplemented with 20 mM acetate and kanamycin for 6 hours. The Tn5 mutant pool was then pelleted and transferred to nitrogen-fixing medium with 20 mM acetate and kanamycin overnight. Metronidazole was then added to the Tn5 mutant pool and the culture was allowed to incubate at 30°C for 8 hours. The Tn5 mutant pool was then washed with minimal medium twice and plated on minimal medium agar with 10 mM succinate as a carbon source supplemented with kanamycin. Roughly 200 individual clones were isolated from this selection and screened for their ability to grow under nitrogen-fixing conditions compared to the parent strain (Δ*fixC\**) by patching each clone on minimal medium agar with 20 mM acetate and kanamycin and nitrogen-fixing medium supplemented with 20 mM acetate and kanamycin.

### Inverse PCR

Transposon mutants were grown in liquid minimal medium with 20 mM acetate to stationary phase, and genomic DNA was purified using a Yeast/Bact Genomic DNA purification kit (QIAGEN, Hilden, Germany). 1 μg of genomic DNA was digested with the restriction enzyme AatII overnight at 37°C to generate fragments of genomic DNA which were on average 1,500 bp (New England Biolabs). The digestion product was then treated with Antarctic phosphatase for 1 hour at 37°C (New England Biolabs). PCR products were purified using the Zymogen Clean and Concentrator PCR-cleanup kit (Zymo Research, Irvine, CA), and the recovered DNA fragments were ligated together to form closed circular DNA using T4 DNA ligase (New England Biolabs). The library of circular DNA fragments was then used as a PCR template with forward and reverse primers specific to the transposable element and amplified using Phusion High-Fidelity DNA Polymerase (New England Biolabs). PCR products were separated by electrophoresis on a 1% agarose gel and purified using the Zymoclean Gel DNA recovery Kit (Zymo Research, Irvine, CA). The purified DNA fragments were then submitted for Sanger sequencing (GENEWIZ, South Plainfield, NJ) using primers specific to the plasmid DNA next to either of the Tn5 mosaic ends (Tn5 SeqF: 5’ – GAGTCAGCAACACCTTCTTCACGAGG – 3’ and Tn5 SeqR: 5’ – GGACAACAAGCCAGGGATGTAACGC – 3’).

### Gas chromatography & protein concentration

H_2_ was quantified using a Shimadzu GC-2014 gas chromatogram equipped with a thermal conductivity detector and a 60/80 molecular sieve 5Å column (6ft x 1/8in.; Supelco). H_2_ standards were measured in technical triplicate and the relationship between the moles of H_2_ in each standard and the area under the detected hydrogen peak was fitted with a linear equation (R^2^ = 0.9976). Samples of headspace taken from growing cultures were measured in biological triplicate and technical duplicate. To measure the H_2_ concentration from growing cultures, the headspace was sampled once the culture reached an optical density of 0.4-0.55 at 660 nm. Cultures were vortexed briefly, then allowed to settle before sampling. H_2_ production in each sample was normalized to the optical density (660 nm) of the culture at the time that the headspace was sampled.

### Protein sequence analysis

Protein sequences and genomic regions were collected from the Joint Genome Institute - Integrated Microbial Genomes and Microbiomes (JGI-IMG/M) database. Specific protein-protein alignments were generated using the Constraint Based Alignment Tool (COBALT, NCBI) using the default parameters. Protein domains were identified using InterPro v.86.0 using the default parameters (26). Homologs of *korAB* were identified using JGI/IMG-M using *R. palustris* BisB5 KorA as a bait sequence (WP_011501953.1). Candidate homologs of *korA* had greater than 80% amino acid identity to *korA* and were adjacent to genes involved with anaerobic benzoate or 4-hydroxybenzoate degradation. AadN (WP_011156245) was used as a bait sequence to identify homologs among the selected *R. palustris* strains. Candidate homologs of *aadN* had greater than 90% amino acid identity and were adjacent to genes involved with anaerobic benzoate or 4-hydroxybenzoate degradation.

### Statistical Analysis

Doubling times of different strains in each growth experiment were compared using ANOVA to test the null hypothesis that the strains come from the same population (P_ANOVA_ < 0.001). Welch’s t-test was used to compare the mean doubling times of individual strains. Similarly, normalized H_2_ accumulation was compared using ANOVA followed by a Welch’s t-test. All statistical analyses were performed in R version 4.1.1.

## Data availability

Genome sequencing data for *R. palustris* Δ*fixC\** has been deposited in the NCBI Sequence Read Archive in BioProject PRJNA858464. RNA-seq data for comparative transcriptomics of *R. palustris* and *R. palustris* Δ*fixC\** has been deposited on the NCBI Gene Expression Omnibus in BioProject PRJNA858255. Homologs of *korAB* were identified using JGI/IMG-M using *R. palustris* BisB5 *korA* as a bait sequence (WP_011501953.1). AadN (WP_011156245) was used as a bait sequence to identify homologs among the selected *R. palustris* strains.

## Acknowledgements

We thank Jack Reddan and Nicholas Haas for their help with strain construction. This work was supported by award DE-SC0020252 from the U.S. Department of Energy, Office of Science, Basic Energy Sciences, Physical Biosciences program to K.R.F.

