## Supplemental figures and tables for "Selecting a new electron transfer pathway for nitrogen fixation uncovers an electron bifurcating-like enzyme involved in anaerobic aromatic compound degradation"

**Supplementary Figures & Tables**

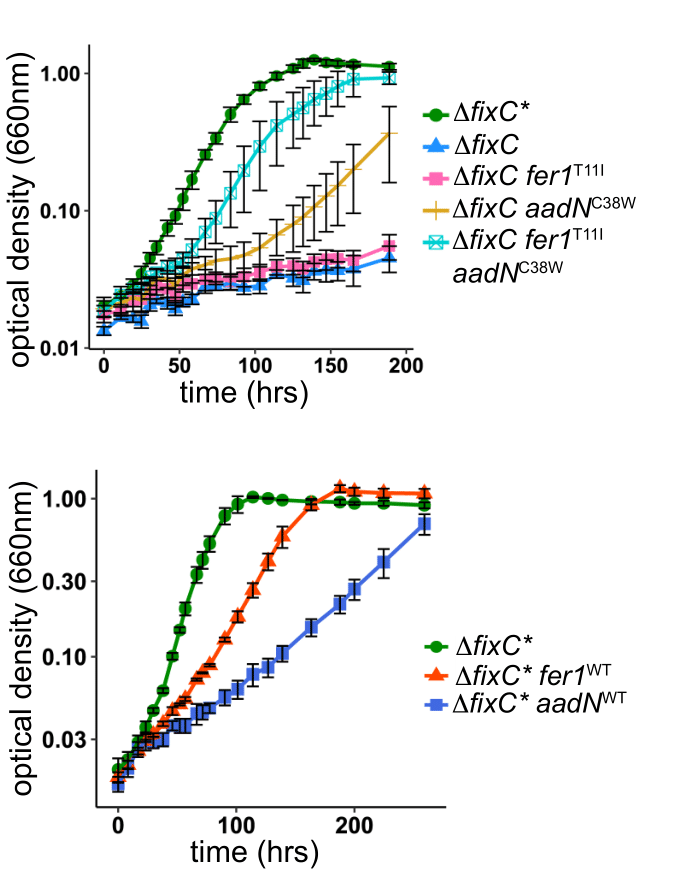

**Figure S1. Repairing mutations in *fer1* or *aadN* in *R. palustris*** Δ***fixC** impairs growth under nitrogen-fixing conditions.** Growth of *R. palustris* Δ*fixC** (Δ*fixC**), *R. palustris* Δ*fixC** with the T11I mutation in *fer1* repaired (Δ*fixC** *fer1*^WT^), and *R. palustris* Δ*fixC** with the C38W mutation in *aadN* repaired (Δ*fixC** *aadN*^WT^) in minimal medium lacking ammonium sulfate (nitrogen-fixing) with 20 mM acetate provided as a carbon source. Data are the average of three biological replicates, and error bars represent one standard deviation.

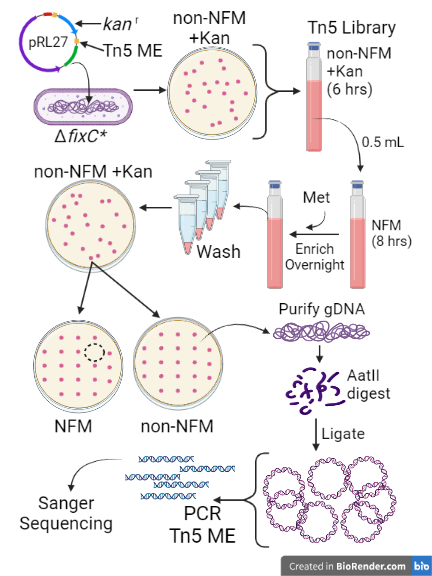
**Figure S2. Workflow for transposon mutagenesis with metronidazole enrichment.** **NFM**, Nitrogen-fixing medium which is minimal medium lacking ammonium sulfate and 20 mM acetate provided as a carbon source; **non-NFM**, minimal medium with 20 mM acetate provided as a carbon source; **Met**, metronidazole; **Kan**, kanamycin; Tn5 ME, Tn5 mosaic end. Incubation times are shown in parentheses.

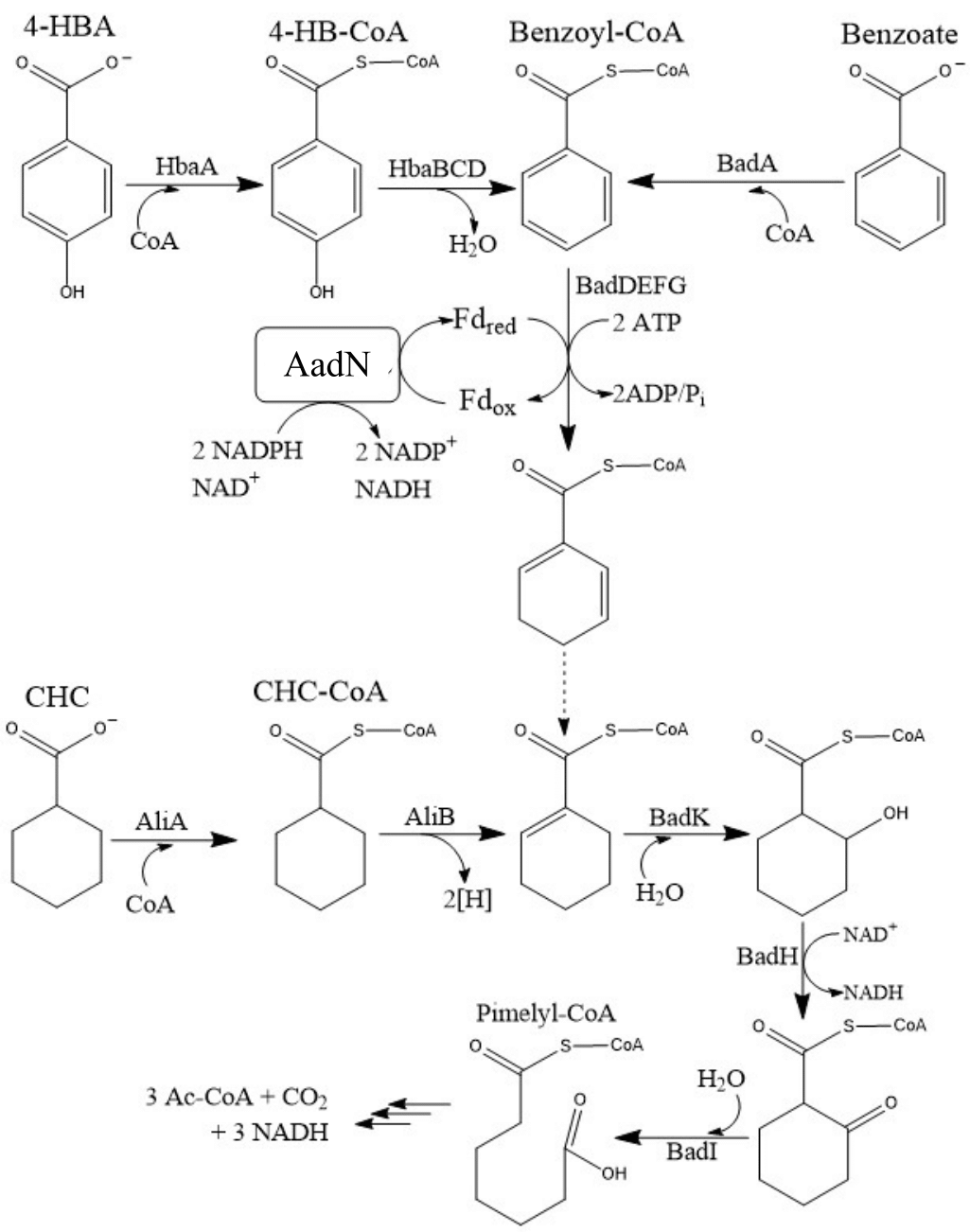
**Figure S3. Degradation pathway of Benzoate, 4-Hydroxybenzoate (4-HBA), and cyclohexane carboxylate (CHC) in *R. palustris* incorporating the proposed activity of AadN.** The activity of AadN is inferred based on its similarity to *Pf*NfnI. The reaction marked by the dotted line represents the uncertainty of whether benzoyl-CoA reductase (BadDEFG) catalyzes a second reduction of the ring structure after the initial de-aromatization step.

**Table S1**. Changes in transcript abundance of genes encoding Fd-reducing enzymes in *R. palustris* Δ*fixC**

| Locus Tag | Gene name | Annotation | Log_2_FC^a^ (Δ*fixC**/WT) | p-value |
| --- | --- | --- | --- | --- |
| RPA0678 | *aadN* | sulfide dehydrogenase, possible glutamate synthase | -0.3 | 0.0067 |
| RPA1228 |  | pyruvate:Fd oxidoreductase | 1.8 | <0.0001 |
| RPA1578 | *cyaA* | Fd:NADP^+^ oxidoreductase | -0.4 | <0.0001 |
| RPA3195 |  | pyruvate:Fd oxidoreductase | 1.5 | <0.0001 |
| RPA3710 |  | Fd:nitrite reductase | -0.8 | <0.0001 |
| RPA4721 |  | pyruvate:Fd oxidoreductase | -0.5 | <0.0001 |
| RPA4722 |  | NAD^+^-dependent Fd:NADPH oxidoreductase | -1.1 | <0.0001 |

^a^ FC: Fold change comparing *R. palustris* Δ*fixC** to wild-type *R. palustris* (WT). “-“ indicates a lower abundance in *R. palustris* Δ*fixC** compared to wild-type.

**Table S2.** All strains, plasmids, and primers used in this study.

| **Strain, plasmid, or primer** | **Genotype or primer sequence (5’ – 3’)** | **Reference** |
| --- | --- | --- |
| ***R. palustris* strains** |  |  |
| CGA753 (wild-type) | CGA009 with an in-frame deletion of *vnfH* and *anfH* | (1) |
| Δ*fixC* | CGA753 with an in-frame deletion of *fixC* | (2) |
| Δ*fixC** | Suppressor strain derived from Δ*fixC*. See Table 1 for genotype | This study |
| Δ*fixC** Δ*fixA* | Δ*fixC** with an in-frame deletion of *fixA* | This study |
| Δ*fixC** Δ*fer1* | Δ*fixC** with an in-frame deletion of *fer1* | This study |
| Δ*fixC** Δ*fer1* Δ*fldA* | Δ*fixC** with an in-frame deletion of *fer1, fldA* | This study |
| Δ*fixC** Δ*fer1* Δ*fldA* Δ*ferN* | Δ*fixC** with an in-frame deletion of *fer1, fldA, ferN* | This study |
| Δ*fixC** *aadN*::Tn5 | Δ*fixC** with a Tn5 insertion in codon 38 of *aadN* in the same orientation as the coding sequence | This study |
| Δ*fixC* *fer1*^T11I^ | Δ*fixC* with the Δ*fixC** allele of *fer1* | This study |
| Δ*fixC aadN*^C38W^ | Δ*fixC* with the Δ*fixC** allele of *aadN* | This study |
| Δ*fixC* *bchN*^S63P^ | Δ*fixC* with the Δ*fixC** allele of *bchN* | This study |
| Δ*fixC hupJ*^T76A^ | Δ*fixC* with the Δ*fixC** allele of *hupJ* | This study |
| Δ*fixC rpa1135*^F224S^ | Δ*fixC* with the Δ*fixC** allele of *rpa1135* | This study |
| Δ*fixC rpa1377*^F250Y^ | Δ*fixC* with the Δ*fixC** allele of *rpa1377* | This study |
| Δ*fixC rpa1470^L138P^* | Δ*fixC* with the Δ*fixC** allele of *rpa1470* | This study |
| Δ*fixC rpa1496*^Q63stop^ | Δ*fixC* with the Δ*fixC** allele of *rpa1496* | This study |
| Δ*fixC rpa1975*^N277S, M356V^ | Δ*fixC* with the Δ*fixC** allele of *rpa1975* | This study |
| Δ*fixC rpa2153*^D230G^ | Δ*fixC* with the Δ*fixC** allele of *rpa2153* | This study |
| Δ*fixC rpa*2193^N152S^ | Δ*fixC* with the Δ*fixC** allele of *rpa2193* | This study |
| Δ*fixC rpa2309*^G317fs^ | Δ*fixC* with the Δ*fixC** allele of *rpa2309* | This study |
| Δ*fixC* *rpa3372*^A74V^ | Δ*fixC* with the Δ*fixC** allele of *rpa3372* | This study |
| Δ*fixC rpa4087*^A68fs^ | Δ*fixC* with the Δ*fixC** allele of *rpa4087* | This study |
| Δ*fixC rpa4534*^T15M^ | Δ*fixC* with the Δ*fixC** allele of *rpa4534* | This study |
| Δ*fixC recQ*^A205T^ | Δ*fixC* with the Δ*fixC** allele of *recQ* | This study |
| Δ*fixC* *fer1*^T11I^ *aadN*^C38W^ | Δ*fixC* with the Δ*fixC** allele of *fer1* and *aadN* | This study |
| Δ*fixC** *fer1*^WT^ | Δ*fixC** with the *fer1*^T11I^ mutation repaired | This study |
| Δ*fixC** *aadN*^WT^ | Δ*fixC** with the *aadN*^C38W^ mutation repaired | This study |
| Δ*aadN* | Mo-only with an in-frame deletion in *aadN* | This study |
| Δ*fixC* Δ*aadN* | Mo-only with an in-frame deletion in *fixC* and *aadN* | This study |
| ***E. coli* strains** |  |  |
| S17-1 | *thi pro hdsR hdsM^+^ recA*; chromosomal insertion of RP4-2 (Tc::Mu Km::Tn7) | (3) |
| DH5-α | *fhuA2* Δ*(argF-lacZ)U169 phoA glnV44 Φ80* Δ*(lacZ)M15 gyrA96 recA1 relA1 endA1 thi-1 hsdR17* | NEB |
| BW20767 | RP4–2-Tc::Mu-1 kan::Tn7 integrant leu-63::IS10 *recA1 zbf-5 creB510 hsdR17 endA1 thi uidA* (∆*MluI*)::*pir*+ | (4) |
| **Plasmids** |  |  |
| pJQ200SK | Gm^r^, *sacB*; mobilizable suicide vector | (5) |
| pRL27_Tn5 | Conjugation-mobilized suicide vector encoding a Kan^r^ transposable element flanked by Tn5 mosaic ends. | (6) |
| pJQ200SK_*fer1*^T11I^ | pJQ200SK with the sequence amplified by the *rpa4631* F and *rpa4631* R primers from Δ*fixC** genomic DNA inserted at the PstI site | This study |
| pJQ200SK_*fer1*^WT^ | pJQ200SK with the sequence amplified by the *rpa4631* F and *rpa4631* R primers from Mo-only genomic DNA inserted at the PstI site | This study |
| pJQ200SK_*aadN*^C38W^ | pJQ200SK with the sequence amplified by the *aadN* usF and *aadN* 1.3kb R primers from Δ*fixC** genomic DNA inserted at the PstI site | This study |
| pJQ200SK_*aadN*^WT^ | pJQ200SK with the sequence amplified by the *aadN* usF and *aadN* 1.3kb R primers from Mo-only genomic DNA inserted at the PstI site | This study |
| pJQ200SK_*bchN*^S63P^ | pJQ200SK with the sequence amplified by the *bchN* F and *bchN* R primers from Δ*fixC** genomic DNA inserted at the PstI site | This study |
| pJQ200SK_*hupJ*^T76A^ | pJQ200SK with the sequence amplified by the *rpa0971* F and *rpa0971* R primers from Δ*fixC** genomic DNA inserted at the PstI site | This study |
| pJQ200SK_*rpa1135*^F224S^ | pJQ200SK with the sequence amplified by the *rpa1135* F and *rpa1135* R primers from Δ*fixC** genomic DNA inserted at the PstI site | This study |
| pJQ200SK_*rpa1377*^F250Y^ | pJQ200SK with the sequence amplified by the *rpa1377* F and *rpa1377* R primers from Δ*fixC** genomic DNA inserted at the PstI site | This study |
| pJQ200SK_*rpa1470*^L138P^ | pJQ200SK with the sequence amplified by the *rpa1470* F and *rpa1470* R primers from Δ*fixC** genomic DNA inserted at the PstI site | This study |
| pJQ200SK_*rpa1496*^Q63stop^ | pJQ200SK with the sequence amplified by the *rpa1496* F and *rpa1496* R primers from Δ*fixC** genomic DNA inserted at the PstI site | This study |
| pJQ200SK_*rpa1975*^N277S, M356V^ | pJQ200SK with the sequence amplified by the *rpa1975* F and *rpa1975* R primers from Δ*fixC** genomic DNA inserted at the PstI site | This study |
| pJQ200SK_*rpa2153*^D230G^ | pJQ200SK with the sequence amplified by the *rpa2153* F and *rpa2153* R primers from Δ*fixC** genomic DNA inserted at the PstI site | This study |
| pJQ200SK_*rpa2193*^N152S^ | pJQ200SK with the sequence amplified by the *rpa2193* F and *rpa2193* R primers from Δ*fixC** genomic DNA inserted at the PstI site | This study |
| pJQ200SK_*rpa2309*^G317fs^ | pJQ200SK with the sequence amplified by the *rpa2309* F and *rpa2309* R primers from Δ*fixC** genomic DNA inserted at the PstI site | This study |
| pJQ200SK_*rpa3372*^A74V^ | pJQ200SK with the sequence amplified by the *rpa3372* F and *rpa3372* R primers from Δ*fixC** genomic DNA inserted at the PstI site | This study |
| pJQ200SK_*rpa4087*^A68fs^ | pJQ200SK with the sequence amplified by the *rpa4087* F and *rpa4087* R primers from Δ*fixC** genomic DNA inserted at the PstI site | This study |
| pJQ200SK_*rpa4534*^T15M^ | pJQ200SK with the sequence amplified by the *rpa4534* F and *rpa4534* R primers from Δ*fixC** genomic DNA inserted at the PstI site | This study |
| pJQ200SK_*recQ*^A205T^ | pJQ200SK with the sequence amplified by the *rpa4826* F and *rpa4826* R primers from Δ*fixC** genomic DNA inserted at the PstI site | This study |
| **Primers** |  |  |
| *aadN* seqF | GCACAAATCAGCCCATCCGTTGATGG | This study |
| *aadN* seqR | GTTCGTTCGCCTTGTCTTCCTCGGTC | This study |
| *aadN* usF | TCACTAAAGGGAACAAAAGCTGGAGTGCAGCGGACCTGCGACG | This study |
| *aadN* usR | CGCGGGCTTACGTCAGCAGCCCGCGGTGCCGCACACTTGCCG | This study |
| *aadN* dsF | GGGCTCGCCGGCAAGTGTGCGGCACCGCGGGCTGCTGACGTAAGC | This study |
| *aadN* dsR | CCGGGGGATCCACTAGTTCTAGAGCGCACCTTCTGCTCGATCACCGAGC | This study |
| *aadN* 1.3kb_R | CCGGGGGATCCACTAGTTCTAGAGCACGCAGGTCGGTGTGATGCAC | This study |
| pJQ200SK Δ*aadN* F | TGCTCGGTGATCGAGCAGAAGGTGCGCTCTAGAACTAGTGGATCCCCCGG | This study |
| pJQ200SK Δ*aadN* R | GCATCGCGTCGCAGGTCCGCTGCACCCTCCAGCTTTTGTTCCCTTTAGTGAGGG | This study |
| pJQ200SK *aadN* WT::C38W F | TCCAGTGCATCACACCGACCTGCGTGCTCTAGAACTAGTGGATCCCCCGG | This study |
| *rpa0971* seqF | GACCGCTGAGGTCGCAGCG | This study |
| *rpa0971* seqR | CATGGCCGCCGCTCCATCC | This study |
| *rpa0971* F | TCACTAAAGGGAACAAAAGCTGGAGCGGTGTGTCCGAGCCTGG | This study |
| *rpa0971* R | CCGGGGGATCCACTAGTTCTAGAGCCCGCCGAGAAACAGAACCGC | This study |
| pJQ200SK *rpa0971* F | CCGCCGAGAAACAGAACCGCATCGCGCTCTAGAACTAGTGGATCCCCCGG | This study |
| pJQ200SK *rpa0971* R | ACCACCGCCAGGCTCGGACACACCGCTCCAGCTTTTGTTCCCTTTAGTGAGGG | This study |
| *rpa1135* seqF | CCAACCACTACCAGATGCAGCTC | This study |
| *rpa1135* F | TCACTAAAGGGAACAAAAGCTGGAGGTGAGCAACATCAAAGGAACGCTGG | This study |
| *rpa1135* R | CCGGGGGATCCACTAGTTCTAGAGCCTGATGGAAGGCTTCGAGCGTCGAG | This study |
| pJQ200SK *rpa1135* F | GACGCCGAACTCGACCGGATCATTCGCTCTAGAACTAGTGGATCCCCCGG | This study |
| pJQ200SK *rpa1135* R | GATTTCGTCGGCGGCGATATGGTAGCTCCAGCTTTTGTTCCCTTTAGTGAGGG | This study |
| *rpa1377* seqF | CACACCACGTCGCCGAGCAC | This study |
| *rpa1377* seqR | GCCGCATCCTGCTGGTGG | This study |
| *rpa1377* F | TCACTAAAGGGAACAAAAGCTGGAGGCTGGCCGACCTCATCGAAGCTC | This study |
| *rpa1377* R | CCGGGGGATCCACTAGTTCTAGAGCGGAATTCGCGTGAATCTTGCTGGCC | This study |
| pJQ200SK *rpa1377* F | GGCCAGCAAGATTCACGCGAATTCCGCTCTAGAACTAGTGGATCCCCCGG | This study |
| pJQ200SK *rpa1377* R | CGGAGCTTCGATGAGGTCGGCCAGCCTCCAGCTTTTGTTCCCTTTAGTGAGGG | This study |
| *rpa1470* F | TCACTAAAGGGAACAAAAGCTGGAGCCCTGCTCGACGTTCGTGATCTCAC | This study |
| *rpa1470* R | CCGGGGGATCCACTAGTTCTAGAGCCTCGGAGCTTCCCTGCTCGACGATC | This study |
| pJQ200SK *rpa1470* F | GATCGTCGAGCAGGGAAGCTCCGAGGCTCTAGAACTAGTGGATCCCCCGG | This study |
| pJQ200SK *rpa1470* R | GTGAGATCACGAACGTCGAGCAGGGCTCCAGCTTTTGTTCCCTTTAGTGAGGG | This study |
| *rpa1496* seqF | CCCACTCTGCCTCCGCTG | This study |
| *rpa1496* seqR | GTAGCCTTCGCTGTGGCGGTAG | This study |
| *rpa1496* F | TCACTAAAGGGAACAAAAGCTGGAGCACCAAGATTCTCGACGGCTACGGG | This study |
| *rpa1496* R | CCGGGGGATCCACTAGTTCTAGAGCCAGACGAAGTCAGCGACCAGATCGG | This study |
| pJQ200SK *rpa1496* F | CCGATCTGGTCGCTGACTTCGTCTGGCTCTAGAACTAGTGGATCCCCCGG | This study |
| pJQ200SK *rpa1496* R | CCCGTAGCCGTCGAGAATCTTGGTGCTCCAGCTTTTGTTCCCTTTAGTGAGGG | This study |
| *bchN* seqF | CATGTCACAGGTTGTTCGACCGC | This study |
| *bchN* F | TCACTAAAGGGAACAAAAGCTGGAGGATGCTCTACACGCCCGAAGAGC | This study |
| *bchN* R | CCGGGGGATCCACTAGTTCTAGAGCGCGAACAATTCAGCGAGATCGGCC | This study |
| pJQ200SK *bchN* F | CGGCCGATCTCGCTGAATTGTTCGCGCTCTAGAACTAGTGGATCCCCCGG | This study |
| pJQ200SK *bchN* R | CCGCTCTTCGGGCGTGTAGAGCATCCTCCAGCTTTTGTTCCCTTTAGTGAGGG | This study |
| *rpa1975* seqF | GACATCTATCCGGCGCTGGAAAAGG | This study |
| *rpa1975* F | TCACTAAAGGGAACAAAAGCTGGAGCGTCGTGACTTTCTGAAAGTGTCAGC | This study |
| *rpa1975* R | CCGGGGGATCCACTAGTTCTAGAGCGATGCAGGGCTTCTGGTTCGTGATC | This study |
| pJQ200SK *rpa1975* F | GATCACGAACCAGAAGCCCTGCATCGCTCTAGAACTAGTGGATCCCCCGG | This study |
| pJQ200SK *rpa1975* R | GCTGACACTTTCAGAAAGTCACGACGCTCCAGCTTTTGTTCCCTTTAGTGAGGG | This study |
| *rpa2153* seqF | GTGAGCCGATCGGCGTCGTC | This study |
| *rpa2153* seqR | CCTTGGTCTTCTCGGCGACG | This study |
| *rpa2153* F | TCACTAAAGGGAACAAAAGCTGGAGGCCGAAGGTGTCGAGACCGAAAAGC | This study |
| *rpa2153* R | CCGGGGGATCCACTAGTTCTAGAGCGATCGCAAAGGCCGTGAACTCAGAG | This study |
| pJQ200SK *rpa2153* F | CTCTGAGTTCACGGCCTTTGCGATCGCTCTAGAACTAGTGGATCCCCCGG | This study |
| pJQ200SK *rpa2153* R | GCTTTTCGGTCTCGACACCTTCGGCCTCCAGCTTTTGTTCCCTTTAGTGAGGG | This study |
| *rpa2193* seqR | GACACCATCAACACCATCAAGCAGG | This study |
| *rpa2193* F | TCACTAAAGGGAACAAAAGCTGGAGGGACTTCCTGGAGCGGGGGTTGATC | This study |
| *rpa2193* R | CCGGGGGATCCACTAGTTCTAGAGCGTCTTTCAGGGGGCGGAAGGCTTGG | This study |
| pJQ200SK *rpa2193* F | CCAAGCCTTCCGCCCCCTGAAAGACGCTCTAGAACTAGTGGATCCCCCGG | This study |
| pJQ200SK *rpa2193* R | GATCAACCCCCGCTCCAGGAAGTCCCTCCAGCTTTTGTTCCCTTTAGTGAGGG | This study |
| *rpa2309* seqF | CGGGTTCGTCACTTTCATCGCC | This study |
| *rpa2309* seqR | CATGCCGTCTGTCCTCTCGC | This study |
| *rpa2309* F | TCACTAAAGGGAACAAAAGCTGGAGGTACGCTCGACACTGCCATGGTTCG | This study |
| *rpa2309* R | CCGGGGGATCCACTAGTTCTAGAGCGACGCTTCGATGCGTTCGCCATC | This study |
| pJQ200SK *rpa2309* F | CGGATGGCGAACGCATCGAAGCGACGCTCTAGAACTAGTGGATCCCCCGG | This study |
| pJQ200SK *rpa2309* R | CGAACCATGGCAGTGTCGAGCGTACCTCCAGCTTTTGTTCCCTTTAGTGAGGG | This study |
| *rpa3372* seqF | GCCGTCCCGCGATCCCATTG | This study |
| *rpa3372* seqR | CAGCGCGACAATTTTTCCAGCTC | This study |
| *rpa3372* F | TCACTAAAGGGAACAAAAGCTGGAGGCTCGTTGCCGTTTCGATGCTGGCC | This study |
| *rpa3372* R | CCGGGGGATCCACTAGTTCTAGAGCCAGTATTTGGCAACCGCCGGACCACC | This study |
| pJQ200SK *rpa3372* F | GGCCAGCATCGAAACGGCAACGAGCGCTCTAGAACTAGTGGATCCCCCGG | This study |
| pJQ200SK *rpa3372* R | GGTGGTCCGGCGGTTGCCAAATACTGCTCCAGCTTTTGTTCCCTTTAGTGAGGG | This study |
| *rpa4087* seqF | CCGTTCAAACGTCCGCCGC | This study |
| *rpa4087* seqR | CGCGCAGCAGAGACCCAAC | This study |
| *rpa4087* F | TCACTAAAGGGAACAAAAGCTGGAGCAGCTCGATTCGATCCAGAGAAAGC | This study |
| *rpa4087* R | CCGGGGGATCCACTAGTTCTAGAGCGAAACCTTGCAACATCTCGTCGATC | This study |
| pJQ200SK *rpa4087* F | GATCGACGAGATGTTGCAAGGTTTCGCTCTAGAACTAGTGGATCCCCCGG | This study |
| pJQ200SK *rpa4087* R | GCTTTCTCTGGATCGAATCGAGCTGCTCCAGCTTTTGTTCCCTTTAGTGAGGG | This study |
| *rpa4534* seqR | GCCATTGTGGTTGCCAGCAGC | This study |
| *rpa4534* F | TCACTAAAGGGAACAAAAGCTGGAGGGTCAGCAACCTGTTTCTGTTCGTTG | This study |
| *rpa4534* R | CCGGGGGATCCACTAGTTCTAGAGCCATTGTCGATAATGGTGGTGCCCTTG | This study |
| pJQ200SK *rpa4534* F | CAAGGGCACCACCATTATCGACAATGGCTCTAGAACTAGTGGATCCCCCGG | This study |
| pJQ200SK *rpa4534* R | CAACGAACAGAAACAGGTTGCTGACCCTCCAGCTTTTGTTCCCTTTAGTGAGGG | This study |
| *rpa4631* seqF | CTTACACCGGCACAGCCCC | This study |
| *rpa4631* F | TCACTAAAGGGAACAAAAGCTGGAGGCGTCTTGCAGGAGCAGGAATTCG | This study |
| *rpa4631* R | CCGGGGGATCCACTAGAGCCCGTGCTCGGCTTCGGAGATC | This study |
| pJQ200SK *rpa4631* F | CGCTGATCTCCGAAGCCGAGCACGGGCTCTAGAACTAGTGGATCCCCCGG | This study |
| pJQ200SK *rpa4631* R | TCGAATTCCTGCTCCTGCAAGACGCCTCCAGCTTTTGTTCCCTTTAGTGAGGG | This study |
| *rpa4826* seqF | CCTACGACGAAGCGAACGACATC | This study |
| *rpa4826* | TCACTAAAGGGAACAAAAGCTGGAGGCTGTCGGTTCTCAACCACGTGTTC | This study |
| *rpa4826* | CCGGGGGATCCACTAGTTCTAGAGCCCGTAATGATCGAGCTTCTTGTCACCG | This study |
| pJQ200SK *rpa4826* | CGGTGACAAGAAGCTCGATCATTACGGGCTCTAGAACTAGTGGATCCCCCGG | This study |
| pJQ200SK *rpa4826* | GAACACGTGGTTGAGAACCGACAGCCTCCAGCTTTTGTTCCCTTTAGTGAGGG | This study |
| Tn5 seqF | GAGTCAGCAACACCTTCTTCACGAGG | This study |
| Tn5 seqR | GGACAACAAGCCAGGGATGTAACGC | This study |
